# A luminal intermediate cell state maintains long-term prostate homeostasis and contributes to tumorigenesis

**DOI:** 10.1101/2023.02.24.529762

**Authors:** Fu Luo, Lara F. Tshering, Karis Tutuska, Mariola Szenk, Diana Rubel, James G. Rail, Savanah Russ, Jingxuan Liu, Alice Nemajerova, Gábor Balázsi, Flaminia Talos

## Abstract

Cellular heterogeneity poses tremendous challenges for developing cell-targeted therapies and biomarkers of clinically significant prostate cancer. The origins of this heterogeneity within normal adult and aging tissue remain unknown, leaving cellular states and transcriptional programs that allow expansions of malignant clones unidentified. To define cell states that contribute to early cancer development, we performed clonal analyses and single cell transcriptomics of normal prostate from genetically-engineered mouse models. We uncovered a luminal transcriptional state with a unique “basal-like” Wnt/p63 signaling (*luminal intermediate*, LumI) which contributes to the maintenance of long-term prostate homeostasis. Moreover, LumI cells greatly expand during early stages of tumorigenesis in several mouse models of prostate cancer. Genetic ablation of p63 *in vivo* in luminal cells reduced the formation of aggressive clones in mouse prostate tumor models. Finally, the LumI cells and Wnt signaling appear to significantly increase in human aging prostate and prostate cancer samples, highlighting the importance of this hybrid cell state for human pathologies with potential translational impact.

## Introduction

Age-related prostate diseases are among the most frequently encountered conditions in the male population and age remains a major risk for prostate cancer (PCa) development. Understanding the cellular origins of prostate hyperplasia and dysplasia and finding novel ways to halt progression towards malignancy remain fundamental challenges of PCa treatment, prevention, and patient stratification.

In general, tissue homeostasis is maintained by constant renewal and differentiation of tissue stem cells. Consequently, stem cell deregulation underlies many diseased conditions, including tumor formation (Blanpain and Simons, 2013). In the adult prostate, several stem/progenitor cell populations have been proposed to replenish lost cells, such as during androgen-dependent organ regeneration. The same cells could also serve as potential cells of origin of prostate cancer (Toivanen and Shen, 2017). Mouse studies have revealed multipotent progenitors active in the basal layer during development (Ousset et al., 2012; Wuidart et al., 2016). During neonatal development when androgen levels are low, a transient population of bipotent luminal cells has also been described (Shibata et al., 2020). In adult, however, the homeostasis relies on largely unipotent lineage committed basal and luminal progenitors, with some evidence for rare bipotent basal progenitors (Choi et al., 2012; Ousset et al., 2012; Wang et al., 2009; Wang et al., 2014; Wuidart et al., 2016; Xie et al., 2017). Nonetheless, the functional progenitors that maintain the homeostasis of androgen-intact normal prostate are not well described, and the deregulations that enable their clonal expansion in tumorigenesis are not well understood.

Intratumor heterogeneity and lineage plasticity are frequently encountered in prostate cancer and pose major diagnostic and treatment challenges (Beltran et al., 2019; Haffner et al., 2021). Prostate cancer is marked by multiple distinct cellular subpopulations of different molecular background and degree of aggressiveness coexisting within the same tumor with implications for adaptation and emergence of therapy-resistant populations under drug-imposed selection (Haffner et al., 2021). Cellular plasticity due to induction of developmental programs or loss of luminal markers also generate heterogeneity and contribute to therapy resistance in prostate cancer (Beltran et al., 2019; Chan et al., 2022; Le Magnen et al., 2018). To date, extensive profiling has revealed the molecular complexity of prostate cancer, yet, understanding the origins and functional roles of altered cellular states within the diverse subpopulations remains limited.

Both mouse and human prostate contain pseudostratified epithelia composed of: *i)* luminal (AR**^+^**, cytokeratin (CK or Krt) 8/18**^+^**), *ii)* basal (p63**^+^**, CK5**^+^**), *iii)* rare neuroendocrine cells, and *iv)* a minor subpopulation of intermediate cells co-expressing basal and luminal markers reported to belong to two positional classes: “basal intermediate” and “luminal intermediate”(van Leenders et al., 2003). Single cell RNAseq further explored the heterogeneity within these main lineages, revealing underappreciated complexity, hybrid states and cellular plasticity (Crowley and Shen, 2022). Prostate cellular heterogeneity likely reverberates into intratumor heterogeneity and impacts drug resistance. In PCa mouse models resistance to antiandrogen drugs has been shown to be accompanied by a loss of the “luminal character” of androgen receptor (AR)-dependent luminal cells and subsequent transition to AR-independent basal-like or neuroendocrine-like cells (Mu et al., 2017; Zou et al., 2017). Drug resistance in prostate cancer has also been linked to plastic transitions initiated in an epithelial population with mixed luminal-basal phenotype dependent on increased JAK/STAT and FGFR signaling (Chan et al., 2022). Luminal cells with basal features have also been shown to expand in the ventral prostate cancer upon Pten-deficiency (Germanos et al., 2022). Recently, single cell transcriptomics of multiple human prostate cancer samples have highlighted increased levels of “intermediate” cells being present in cancer vs the normal tissue (Chen et al., 2021; Song et al., 2022). Thus far, the precise origin of the hybrid cancer states within the normal prostate cellular heterogeneity remains unknown.

In this study, we coupled single cell transcriptomics with multicolor lineage tracing in normal prostate to delineate the long-term luminal clonal dynamics and provide insights into luminal cellular heterogeneity. We show here that the long-term luminal homeostasis is maintained by multiple clones with various levels of fitness/clonal activity. The clones that become larger long-term appear to be maintained through a luminal transcriptional state harboring basal markers (Luminal Intermediate, **LumI**) and controlled by Wnt/p63 signaling. Moreover, cells with LumI characteristics appear to expand with aging in both mouse and human prostate samples. LumI cells also contribute to clonal growth in several mouse models of prostate cancer and associate with numerous human prostate cancer glands. Specific genetic ablation of p63 from luminal cells reduces clonal expansions in prostate cancer of luminal origin highlighting the important role that the LumI transcriptional program has in tumorigenesis. Our models reveal a continuity of this cell state from normal homeostasis to aging and early cancer with specific alterations at each stage.

## Results

### Slow clonal dynamics and clones with different fitness potential sustain the homeostasis of the prostate luminal layer

To assess the clonal dynamics of adult luminal prostate cells, we exploited a stochastic multicolor Cre reporter, *R26R-Confetti* (Schepers et al., 2012) and the *Nkx3.1-CreERT2* knock-in allele where a tamoxifen-inducible Cre-recombinase cassette is placed under the transcriptional control of the prostate-specific Nkx3.1 promoter (Wang et al., 2009). In intact adult prostate, Nkx3.1 homeobox transcription factor is prostate specific and expressed in all luminal cells and very rare basal cells (Bhatia-Gaur et al., 1999), with a preference of expression for the medial-distal prostate (Crowley et al., 2020). Upon tamoxifen administration, Nkx3.1**^+^** cells stochastically express one of the four Confetti colors (RFP, YFP, GFP and CFP) as a permanent genetic mark transmitted to all progeny of the labeled cell enabling clonal spatial identification.

To this end, we administered low dose of tamoxifen (TAM) to adult *Nkx3.1^CreERT2/+^; R26R- Confetti/+* (***N***) mice at 2 months of age when prostate is fully developed and the effects of Nkx3.1 allelic heterozygosity are not yet overt (Bhatia-Gaur et al., 1999; Bowen et al., 2020) (**Fig. 1A**). We used a low dose of tamoxifen to ensure sparse labeling of Nkx3.1**^+^**cells and avoid labeling neighboring cells with the same fluorescent color. After tamoxifen induction, we harvested prostate tissue at various time points of lineage-tracing and performed direct visualization of spatially separated Confetti**^+^** clones by confocal microscopy. At 1 week almost all labeled cells were solitary (clusters of 1 cell) (**Fig. 1B**). The labeling efficiency within the luminal layer was 3.38% +/- 0.35. At next time points, the labeled population is dominated by the presence of single cells consistent with a slow commitment to expand (Wang et al., 2013; Wuidart et al., 2016) (**Fig. 1C-F**). However, by 4 weeks, and more visible by 8 and 12 weeks, a minority of initially labeled single cells grew into larger clusters (>10 cells), which are of clonal origin, based on the unique Confetti color and histological separation (**Fig. 1E-F**, arrows). The larger clones appeared of relatively constant size across the later time points analyzed presumably due to slow cell division or loss of clonal cells. Clonal expansions are relatively rare and demonstrate the delicate control of homeostatic cell replacement in adult prostate tissue.

**Figure 1.**
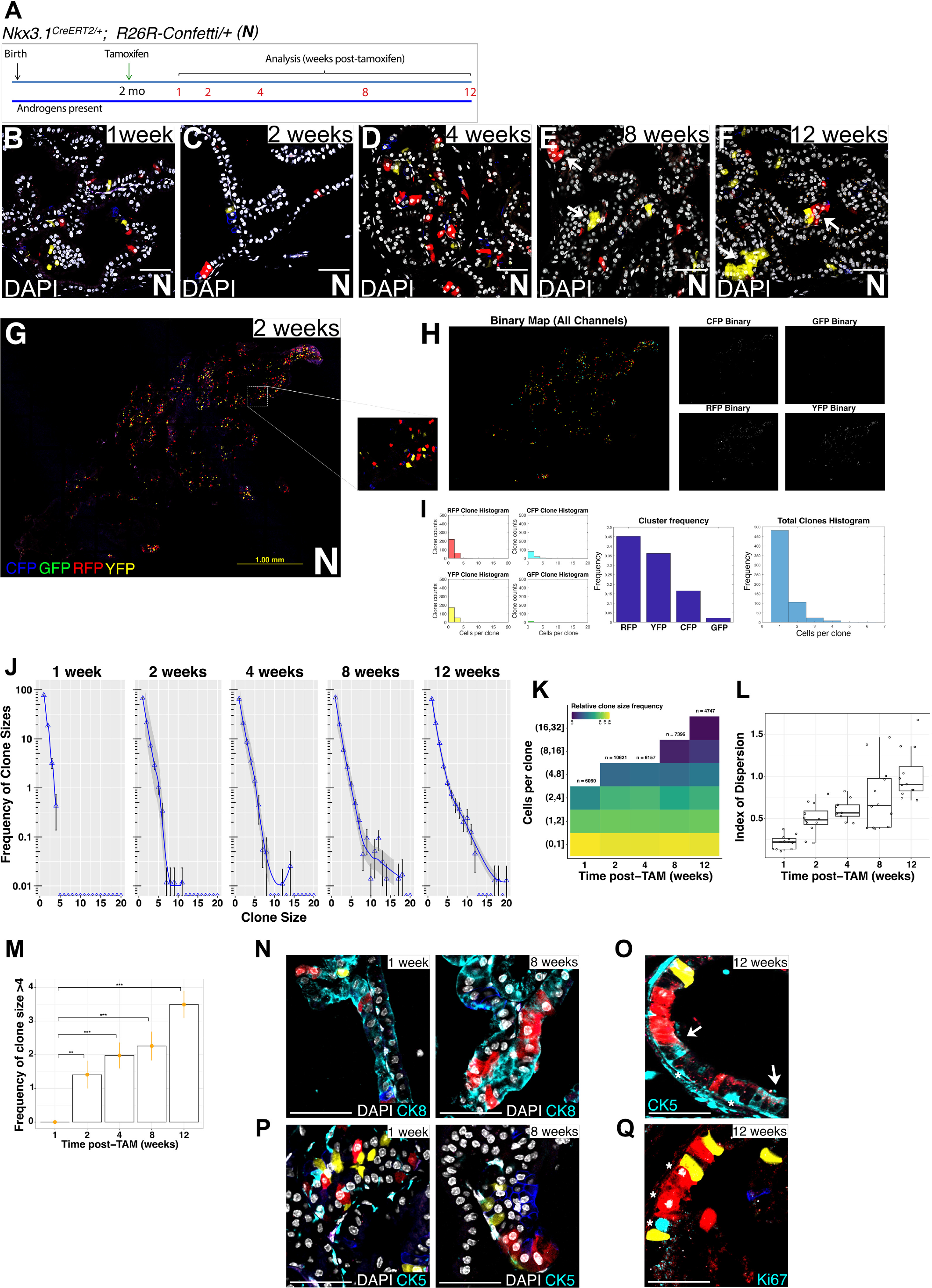
Slow clonal dynamics and clones with different fitness potential sustain prostate luminal homeostasis (**A**) Time course of lineage-tracing experiments with the Confetti reporter in *Nkx3.1^CreERT2/+^* (*N*) mice. (**B-F**) Representative image of clonal distribution in the N-Confetti anterior prostate (AP) at 1, 2, 4, 8 and 12 weeks post-TAM. 40x images are shown, white arrows point to larger clones at late timepoints, scale bars represent 50 μm. (**G-I**) Workflow of clonal quantification by automated image processing. (**G**) Representative image of prostate clonal distribution in *N-Confetti* at 2 weeks post-TAM, showing a 20x scan of an entire section of an AP lobe, with an individual tile shown in inset. (**H**) The image in G was processed through a script developed with the Image Processing toolbox in MatLab. The background subtraction and binarization steps are shown. (**I**) Clonal quantification of the processed image, showing cells/individual clones for each channel (left), total number of clones/each channel/entire section (middle), and distribution of cells/clones for the entire section (right). (**J**) Frequency of clonal size distribution plot, data extracted from clonal quantification of merged tile scan images of entire prostate sections. Points show average of percentages of clones in a given size per slide per timepoint; the blue lines are fitted with the “*geom_smooth*” function in ggplot2 using a locally weighted method (LOESS) of regression and the grey bands are the 95% confidence interval of the regression lines. 24360 clonal events across all time points were included. (**K**) Heatmap representation of clonal size distribution comparing densities of clones of various sizes at indicated timepoints post-TAM. (**L**) Index of dispersion of clonal sizes calculated as the ratios of the variance to the mean of each individual section. (**M**) Percentage of clones with clonal sizes >4 at the indicated time points post-TAM. Percentages are calculated by individual sections. Mean (dot) and SEM (error bar) of percentages are shown. (**N**) Immunofluorescence analysis of luminal marker (CK8) in Confetti clones at 1 week and 8 weeks post-TAM. A general CK8-positivity of Confetti+ cells at both time points is observed. (**O**) Luminal intermediate cells are identified in the luminal layer using a CK5 antibody with high sensitivity. Arrows show luminal Confetti**^+^** cells expressing low levels of CK5. (**P**) Immunofluorescence analysis of basal cells (CK5^high^) in Confetti clones at 1 week and 8 weeks post-TAM. Note the lack of high CK5**^+^** Confetti+ cells in clones in this analysis with a regular CK5 antibody that recognizes only high levels of CK5. (**Q**) Immunofluorescence analysis of Ki67**^+^** staining. Asterisks show proliferative luminal Confetti**^+^** cells with positive Ki67**^+^** staining.

For a precise quantification of clonal dynamics, we performed systematic confocal tile scanning of entire prostate sections collected systematically throughout the anterior (AP) prostate lobes of 4-5 ***N*** mice per time point. First, we acquired images of entire prostate sections (**Fig. 1G**) by confocal tile scanning and merging of the acquired “tiles.” Next, we used an automated script based on a previously published method (Lee et al., 2014) that detects individual regions of fluorescence in each channel (**Fig. 1H**) and counts color clusters and DAPI-positive nuclei (**Fig. 1I**). This allowed a precise quantification of clonal size distribution and showed the progressive accumulation of larger clones as more of the initially labeled cells enter cell cycle and the loss of clones of 1 cell with increased time of tracing (**Fig. 1J, 1K**, **Suppl. Fig. 1A**, 4 mice/time point, 4747 - 10621 clones/time point). In accord, the overall proportion of Confetti**^+^** cells remains relatively constant across time points, with a significant increase at 12 weeks of tracing, correlating with the presence of larger clones (**Suppl. Fig. 1B**). We also observed that the majority of the labeled cells are represented by single cells across all time points (77.77%+/-1.15) likely representing the quiescent differentiated luminal cell population. Focusing on the growing clones, we demonstrate that both the number and the frequency of larger clones increase with longer tracing time. The diversity of clone sizes measured by the index of dispersion (**Fig. 1L**) and the Gini index (**Suppl. Fig. 1C)** increased with time, indicating a more heterogenous clonal size distribution at later time points due to the expanding size of high fitness clones. The frequency of clones with more than 4 cells (considered as growing clones) steadily increases with time (**Fig. 1M)**, suggesting that more of the labeled single cells become proliferative and initiate clonal expansions.

We performed similar analyses to test the generality of these findings in other prostate lobes. The clonal dynamics were similar in the corresponding dorsolateral prostate (DLP) and ventral prostate (VP) lobes where most labeled cells remained quiescent, but few larger clones appeared by 8 weeks (**Suppl. Fig. 1D**). The prostate tissue of adult ***N*** mice appeared to have normal histology at all time points (**Fig. 1B-F, Suppl. Fig. 1E)** and the prostate AP lobes did not show any significant change in weights throughout the duration of our labeling experiments (**Suppl. Fig. 1F**) in accord with previous studies of *Nkx3.1^CreERT2/+^* mice (Wang et al., 2009). We observed similar clonal dynamics in middle aged ***N*** mice (13-16 mo of age) with increased accumulation of larger clones at 8-12 weeks of tracing **(Suppl. Fig. 1G, H)** while the total counts of clones remain constant, similar to the 2 months group (**Suppl. Fig. 1I**).

Given the fact that Nkx3.1 is expressed in luminal cells and rare basal cells, we next asked which initially labeled cells might give rise to clones and where are they located. For this purpose, we manually examined and counted cell types in 36 individual 20x images from 3 individual AP sections per mouse from 4 mice at 1 week and 4 mice at 8 weeks post-tamoxifen. As Nkx3.1 expression is largely confined in adult tissue to the luminal layer, at induction, Confetti colors are preferentially expressed in luminal cells while Confetti-labeled basal cells and triangular “basal intermediate” cells are extremely rare (**Suppl. Fig. 1K,** 1 week post-TAM). Moreover, the rare “basal intermediate” cells remained solitary at all timepoints, and we did not find them as part of the larger clones (**Suppl. Fig. 1K**, 8 weeks post-TAM). The initial Confetti labeling efficiency in the luminal layer was also similar across multiple ***N*** mice at 1 week post-tamoxifen indicating that the clonal results in mice collected at later timepoints can be confidently correlated with this initial seeding (**Suppl. Fig. 1L**). In accord with higher Nkx3.1 levels in the medial/distal prostate ducts (Crowley et al., 2020), we did not find Confetti cells and ensuing clones in the periurethral/proximal prostate region (**Suppl. Fig. 1J**).

To confirm the nature of Confetti**^+^** cells, we performed immunofluorescence (IF) staining with CK5 antibody specific for basal, and CK8 antibody specific for luminal cells. This confirmed the positional assessment of labeled cells as luminal, basal or basal intermediate. All the cells in clones were positive for luminal markers (CK8, **Fig. 1N**) but rare luminal cells in larger clones showed a weak luminal CK5 staining when stained with a highly sensitive CK5 antibody (**Fig. 1O**, arrows). These were clearly identifiable as luminal cells due to their typical luminal position and elongated morphology perpendicular per basement membrane and distinct from basal or small triangular “basal intermediate cells” which have higher levels of CK5 and are not part of luminal clones (**Fig. 1O**, asterisk, and **Fig. 1P**, basal layer stained with regular CK5 antibody). These CK5- low luminal clonal cells (**Fig. 1O**) might encompass the rare “luminal intermediate cells” described previously by histopathological studies (van Leenders et al., 2003). We also show Ki67**^+^** luminal nuclei in larger clones (**Fig. 1Q,** asterisks), a rare find given the fact that the luminal layer is only slowly proliferating, and our lineage tracing is homeostatic. It further shows that luminal cells acquire proliferating capacity to generate larger clones.

In sum, the results of our labeling strategy delineate the clonal expansions originating in the luminal cells and contributing to homeostatic maintenance of the luminal layer. Our results confirm that the luminal layer is maintained by unipotent progenitors that can only generate luminal cells in adult and middle age homeostasis. The long-term luminal homeostasis is maintained by multiple clones with various levels of fitness/clonal expansion activity.

### scRNAseq of Nkx3.1-lineage traced cells reveals cellular heterogeneity within the luminal layer

To understand what luminal subtypes sustain the growth of the larger clones emerging with longer time of lineage tracing, we performed scRNAseq analyses of FACS sorted Confetti**^+^** cells from AP lobes of ***N*** mice. To ensure that Confetti labeling will encompass all luminal subtypes, we maximized the tamoxifen dosage and pooled 4 mice/time point to obtain sufficient labeled cells at two time points, as follows. First, we sequenced cells at **1 week** post-tamoxifen, which includes all initially labeled cells, giving a snap-shot of all luminal cells at homeostasis. Second, we sequenced cells at **8 weeks** of tracing, which includes the progeny of the cells from 1 week that grew into larger clones, the cells at 1 week that did not undergo growth or turnover, and also reflects the cells lost due to turnover. Upon elimination from downstream analyses of the small clusters of contaminating non-epithelial, non-prostatic, and dying cells (**Suppl. Fig. 2A, B**), we found 7 separate epithelial cell clusters present in both 1 and 8 weeks (**Fig. 2A**), each identifiable by specific markers (**Fig. 2B, Suppl. Fig. 2C, Suppl. Table 1** – all markers per cluster, **Suppl. Table 2** – top 5 markers per cluster). While all clusters expressed luminal *Krt8/18 (CK8/18)* and *Ar*, a subset of cells expressed additional basal markers *Krt5* (CK5) and *Trp63* (p63) which we categorized as a “hybrid/intermediate” state and collectively named **LumI** (Intermediate 1 and Intermediate 2 clusters). A separate set of clusters (Secretory 1, Secretory 2, and Secretory 3; collectively named **LumS**) showed high levels of genes involved in prostate specific secretions (including Probasin – *Pbsn*, and *Msmb*) and likely terminally differentiated (**Fig. 2A, 2B**). They might be also implicated in local immune response as they express high levels of *Pigr* (a member of the immunoglobulin superfamily), and Defensins (e.g., *Defb1*, *Defb50*) (**Fig. 2B**). The small **Ki67^+^**cluster reflects the low proliferation rates of adult luminal cells and uniquely expresses factors associated with cell division, *Stmn1* and *Cenpf* (**Fig. 2A-C**). Likely representing the CK4**^+^**cells in invagination tips (Guo et al., 2020), we also detected a small **Krt4^+^**cluster uniquely expressing Krt4 (CK4) and Psca but lacking other basal markers such as CK5 and p63 which are present only in the **LumI** (**Fig. 2A-2C**). Our scRNAseq analysis validates the immunostaining and positional cell type analysis of **Fig. 1**, including the presence of luminal intermediate (CK5**^+^**) cells (**Fig. 1O**), and further refines the luminal cell states of the medial-distal AP prostate.

**Figure 2.**
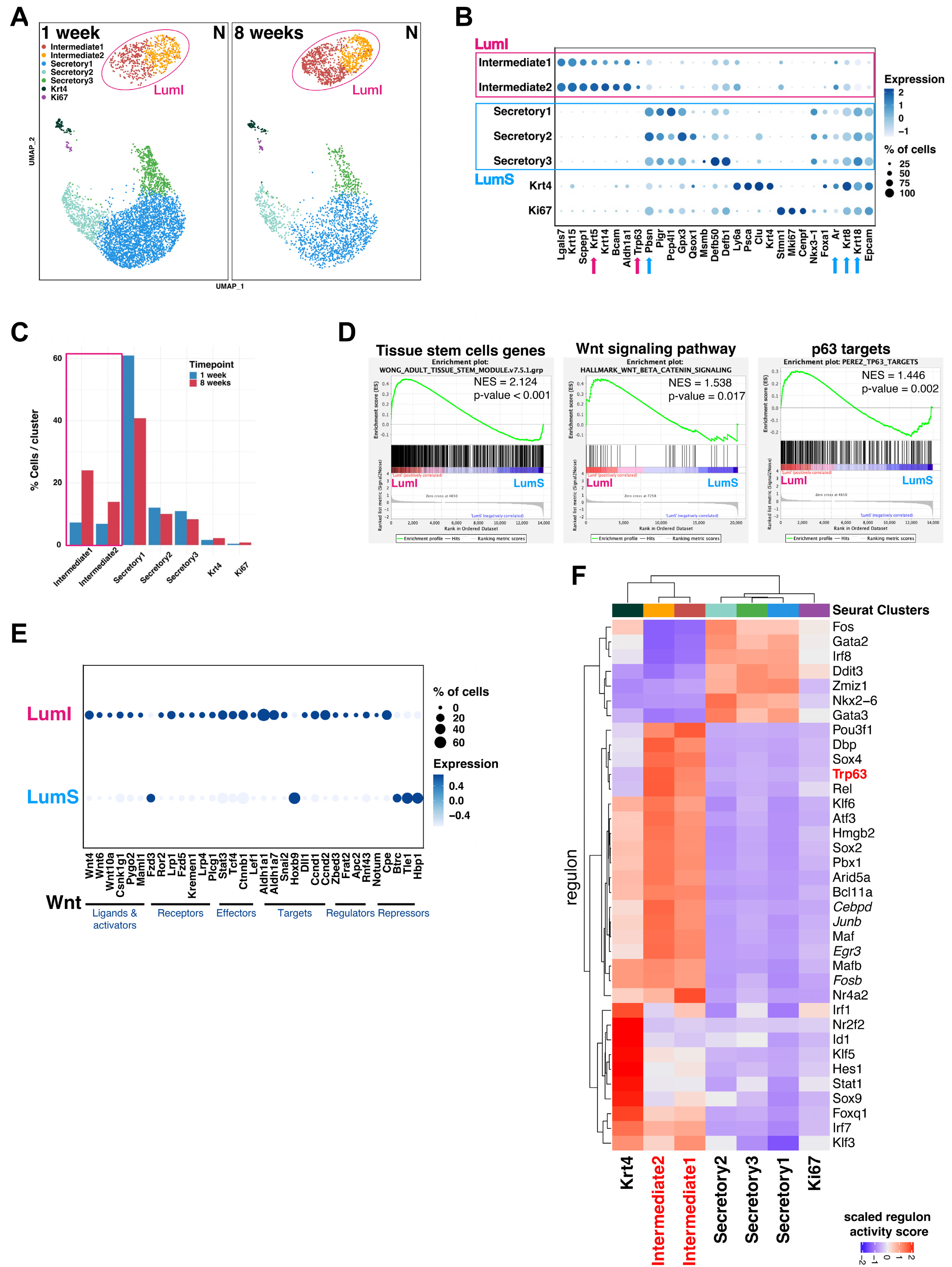
scRNAseq analysis reveals a unique luminal intermediate (LumI) cell state in *N- Confetti* mice. **(A)** Cell type cluster distribution of 1 week and 8 weeks post-TAM *N* samples by UMAP (Seurat) identifying 7 separate epithelial clusters in both samples. LumI cell state is highlighted with red circles. (**B**) Dotplot of selected marker genes in epithelial clusters. (**C**) Percentage of cells in each cluster in the 1 week and 8 weeks samples. Note the increase in the Intermediate1 and Intermediate2 clusters at 8 weeks and the reduction in the Secretory clusters. (**D**) GSEA of adult tissue stem cell markers, canonical Wnt signaling pathway and p63 targets indicate enrichment of the LumI cell state in these three gene sets. For simplicity, the Intermediate1 and Intermediate2 cells were integrated in one Intermediate cluster, while the Secretory1, 2, and 3 were integrated in one Secretory cluster. (**E**) Dotplot of Wnt signaling component genes in intermediate (LumI) and secretory (LumS) clusters. (**F**) Regulon analysis by *SCENIC* implicates *p63* as a master regulator gene of the intermediate clusters.

### The LumI transcriptional state is marked by increased Wnt signaling and p63 pathway activation

Both 1 and 8 weeks cell populations are homeostatic, thus they should contain the entire variety of labeled luminal subtypes. However, the 8 weeks sample includes also the “younger” cells generated during the span of the 8 weeks of tracing by the cells entering proliferation from the initially marked cells at 1 week. We reasoned that the comparison between these two time points should reveal the changes due to clonal expansions and contractions. Indeed, when we compared these samples, the 8 weeks timepoint contained a larger proportion of cells in the LumI clusters at the expense of cell loss from the LumS clusters (**Fig. 2C**). The expansion of the intermediate population during the 8 weeks of tracing coincides with the appearance of larger clones at 8 weeks in our confocal imaging (**Fig. 1B-F**). Thus, the increasing Intermediate population likely represents the newly generated cells in clones and should provide insights into the transcriptional program employed by the luminal cells for clonal growth and maintenance of homeostasis.

We found uniquely enriched pathways in LumI (Intermediate) cells by GSEA (Subramanian et al., 2005), including pathways involved in epithelial proliferation and differentiation: Wnt signaling and mitotic spindle (**Suppl. Fig. 2D**). On the other hand, the LumS (Secretory) clusters are indeed marked by pathways involved in protein secretions, fatty acid metabolism, and oxidative phosphorylation (**Suppl. Fig. 2D**). We confirmed similar LumI activated pathways with *clusterProfiler* (Yu et al., 2012): e.g., positive regulation of cell population proliferation, mitotic cell cycle, positive regulation of cell differentiation (**Suppl. Fig. 2E**). In specific GSEA analyses, we found that LumI cells showed significant enrichment of gene sets associated with adult stem cells (Wong et al., 2008), Wnt canonical signaling pathway and a set of genes upregulated by p63 (Perez et al., 2007) (**Fig. 2D**). Indeed, compared to LumS, the LumI cells have increased Wnt signaling, including higher ligands (e.g., the atypical *Wnt4* and *Wnt6*), receptors (*Lrp1/4*, *Fzd5*), effectors (*Stat3*, *Tcf4*, *Ctnnb1*), target genes (*Aldh1a1*, *Aldh1a7*, *Ccnd2*), modulators (*Zbed3*, *Frat2*), and reduced Wnt repressors (*Tle1*, *Hbp1*) (**Fig. 2E**). Regulon analyses by *SCENIC* (Aibar et al., 2017) revealed specific master regulators (MRs) coordinating these clusters, including an activated p63 response and cell cycle TFs (e.g. *Cebpd*, *Junb*, *Egr3*, *Fosb*) in the LumI (**Fig. 2F**) reflecting their growing phase and implicating p63 in controlling a gene network involved in luminal layer maintenance. Albeit transcriptionally similar, the LumI 8 weeks cells exhibited several significantly enriched pathways compared to LumI cells at 1 week, likely capturing the signaling involved in the LumI expansion: e.g., cellular response to fibroblast growth factor stimulus, response to fibroblast growth factor, endothelial cell chemotaxis, and regulation of keratinocyte differentiation (**Suppl. Fig. 2F**).

In sum, our data suggests that the luminal clonal expansion is maintained by a “luminal-basal” intermediate state activated in luminal cells potentially through coordinated Wnt-p63 signaling and having a progenitor-like activity in clonal expansions.

### LumI cells expand in PCa models

Given their growth program, we asked how LumI cells, and the general luminal heterogeneity are altered during tumorigenesis. For this purpose, we incorporated conditional alleles of Pten deletion (***NP***) and Pten deletion/mutant Kras^G12D^ activation (***NPK***) (Aytes et al., 2013; Aytes et al., 2014) in *Nkx3.1^CreERT2^-Confetti* models and followed similar lineage tracing (**Fig. 3A**). While the NP model is slow progressing, the NPK model is very aggressive, invasive and metastatic (Aytes et al., 2013; Aytes et al., 2014). By 4-5 weeks of tracing, ***NP*** mice developed well delimited and similarly sized tumor clones (**Fig. 3B-D**) corresponding to low grade PIN (**Fig. 3E**) while ***NPK*** mice underwent rapid clonal growth and reorganization leading to a dominant/minor landscape (**Fig. 3F-H**) and high-grade PIN (**Fig. 3I**). At this stage, we collected Confetti**^+^** tumor cells by FACS, and performed scRNAseq. To uncover cell type relationships between the normal and malignant luminal cells, we integrated the scRNAseq data from the ***N***, ***NP*** and ***NPK*** samples. After excluding the small non-epithelial contaminant clusters (**Suppl. Fig. 3A)**, we identified 9 epithelial clusters (**Fig. 3J**) with distinct gene expression (**Suppl. Table 3** - all markers per cluster, **Suppl. Table 4** - top 5 markers per cluster). All clusters were luminal and epithelial (**Suppl. Fig. 3B**, e.g., *Krt8/18**^+^**, EpCam**^+^**, Ar**^+^***), a good representation of human adenocarcinoma. Most clusters were found across all samples and were identifiable using the ***N*** scRNAseq markers (Ki67**^+^**, Krt4^+^, Intermediate 1 and 2, Secretory 1, 2 and 3). Two clusters were mainly associated with ***NP*** and ***NPK***: a Progenitor-like (*Ly6d^+^* and *Tacstd2^+^*, previously reported to mark progenitors (Barros-Silva et al., 2018; Goldstein et al., 2008)), and an EMT (high in EMT markers including Vimentin, *Vim^+^*) cluster (**Fig. 3J, Suppl. Fig. 3B**).

**Figure 3.**
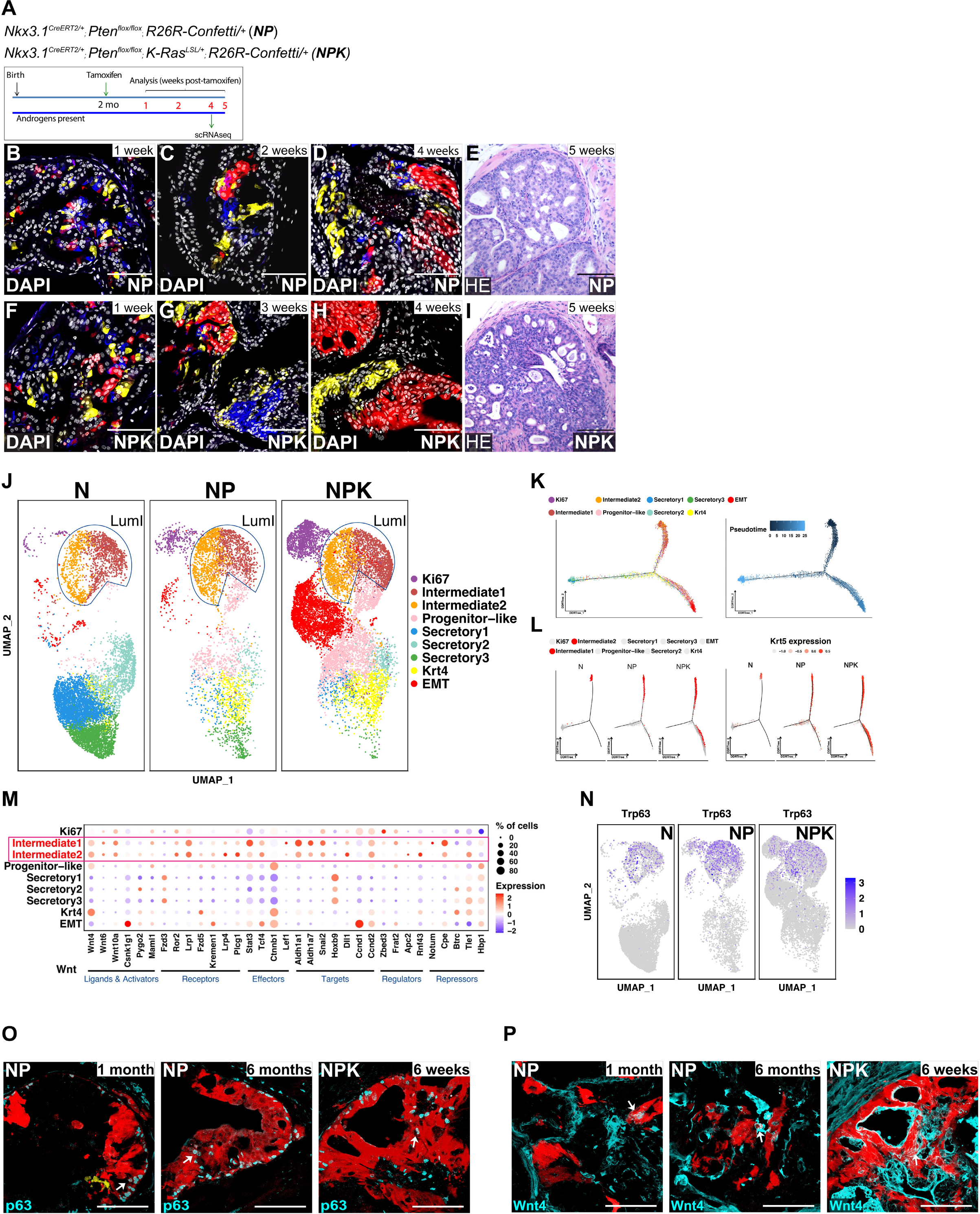
The LumI cell state expands in PCa models **(A)** Time course of lineage-tracing experiments with the Confetti reporter in *NP* and *NPK* mice. (**B- D**) Representative images of clonal distribution in the *NP* anterior prostate at 1, 2 and 5 weeks post-TAM, 40x images are shown, scale bars represent 50 μm. (**E**) Representative H&E staining of *NP* tumors at 5 weeks post-TAM showing features of low-grade PIN. Scale bar represents 100 μm. (**F-H**) Representative images of clonal distribution in the *NPK* anterior prostate at 1, 2 and 4 weeks post-TAM, 40x images are shown, scale bars represent 50 μm. (**I**) Representative H&E staining of *NPK* tumors at 5 weeks post-TAM showing features of high-grade PIN. Scale bar represents 100 μm. (**J**) Cell type cluster distribution of *N*, *NP* and *NPK* Confetti**^+^** samples by UMAP (Seurat) identifying 9 separate epithelial clusters. Transcriptomic *N* features are maintained in the cancer samples. (**K**) Branched trajectory of epithelial cell state transition inferred by Monocle indicating that intermediate cells could serve as cell of origin population and generate the other cell types. Cells are labeled with Seurat clusters on the left panel and *pseudotime* on the right panel. (**L**) Trajectory analysis by Monocle highlighting the intermediate cell states (left) and Krt5**^+^** luminal cells (right) indicating their expansion into multiple branches, including towards the EMT branch. (**M**) Dotplot of selected Wnt pathway genes in the epithelial clusters. Note the increased expression in the Intermediate cell clusters (highlighted in red). (**N**) Feature plot of *Trp63* (p63) expression levels in *N*, *NP* and *NPK* Confetti**^+^**samples. (**O**) Immunofluorescence analysis of p63 expression validating the presence of LumI cells in *NP* and *NPK* Confetti clones. (**P**) The expression of Wnt4 at protein level in LumI cells was validated by immunostaining of *NP* and *NPK* Confetti clones.

In this integrative analysis, the **LumI** state (marked by co-expression of *Krt5* and *Krt8*) emerged as a major contributor to tumor growth (**Fig. 3J, Suppl. Fig. 3C**), while LumS are less represented. Pseudotime trajectory analysis emphasized the distinct origin, expansion and common branching of the LumI cells, and revealed a progression from LumI (top branch) towards LumS and other cancer specific states such as EMT (**Fig. 3K**). In addition, the cells with positive Krt5 (CK5) expression expand similarly as LumI (Intermediate 1&2) along the trajectory, showing the same intermediate state distribution (**Fig. 3L**). Like the ***N*** LumI, the integrated **LumI** cells show increased Wnt signaling (**Fig. 3M**), and enriched adult tissue stem cell, Wnt signaling and p63 targets (**Suppl. Fig. 3D**). p63 also remained as a LumI MR by *SCENIC* (**Suppl. Fig. 3E**). p63 levels increased in cancer both at RNA and protein levels (**Fig. 3N, O**). We also confirmed the presence of Wnt4**^+^** cells in NP and NPK clones (**Fig. 3P**). Of note, both p63 and Wnt4 are present in ***NP*** and ***NPK*** tumors at later time points of lineage tracing (**Fig. 3O, 3P**, 6 months *NP*, 6 weeks *NPK*) indicating that these cells are generated in tumors for a long time and contribute to clonal growth. However, the *NP* and *NPK* LumI cells are likely responding to different signaling as the FGF interactions characteristic for the *N* LumI cells switches to increased Laminin, THBS, EGF, and FN1 ligand-receptor pairs by *CellChat* (Jin et al., 2021) analysis (**Suppl. Fig. 3F**). Thus, the transformed LumI state continues to employ a Wnt-p63 program to engage in malignant proliferations, albeit by engaging additional signaling pathways.

### Specific deletion of p63 in luminal cells reduces the number of growing clones and the clonal size in normal prostate and PCa

To provide insights into the functional role of p63 in *normal* LumI clonal expansions and the effect of perturbing the LumI cells on the general clonal dynamics, we generated a novel mouse model: *Nkx3.1^CreERT2/+^ΔΝp63^flox/flox^ R26R-Confetti*/+ (***Np63***). The transcription factor p63, a member of the p53 family, is a master regulator of stem cell maintenance and differentiation of all epithelial tissues (Botchkarev and Flores, 2014). Its main N-terminal isoforms, TAp63 and ΔΝp63, have pleiotropic roles in tumorigenesis (Su et al., 2013). As indicated by previous studies of p63 isoforms in prostate (Sethi et al., 2015), FACS sorted Confetti**^+^** cells expressed significantly higher levels of the ΔΝp63 isoform compared to almost undetectable levels of TAp63 (**Suppl. Fig. 4A**). *ΔΝp63^flox/flox^* mice (Chakravarti et al., 2014) were recently used to demonstrate the role of ΔΝp63 in promoting tumor growth in a model of lung cancer driven by mutant K-Ras (Napoli et al., 2022). The ***Np63*** prostate shows normal histology (**Suppl. Fig. 4B**) and the lobe weights remained unchanged for the duration of the experiment (**Suppl. Fig. 4C)**. Upon tamoxifen, Nkx3.1**^+^** cells should express one of the Confetti colors and also recombine the p63^flox^ alleles generating *p63- deficient luminal cells*. The Confetti-labeling efficiency was similar to ***N*** mice (**Suppl. Fig. 4D**). Using similar lineage tracing and clonal quantifications (**Fig. 4A**), we observed the same ***N*** dynamics in ***Np63*** mice: while most of the labeled cells remained as single cells, a small subset of larger clones emerged at later timepoints (**Fig. 4B-F**). However, there was a significant loss of clones/section at 8 and 12 weeks in the ***Np63*** mice compared to ***N*** mice (**Fig. 4G**) and this was also demonstrated by the reduction of total Confetti labeled cells at 12 weeks compared to 1 week (**Suppl. Fig. 4D**). The loss of clones is likely due to the increased turnover of a subset of labeled luminal cells. These cells are sensitive to the loss of p63, suggesting that some LumI cells which would otherwise persist and form larger clones are lost. Moreover, the larger size clones appeared at a slower pace in the ***Np63*** reflected by a reduced fold change in average clone size at 2 and 4 weeks post-induction (**Fig. 4H**) and a slower rise of the Gini index during the same time frame (**Fig. 4I**). This suggests that the loss of p63 in luminal cells decreases the clonal fitness in a subset of cells which originally might have had higher fitness. Nevertheless, larger clones do appear by 12 weeks, likely due to heterozygous *ΔΝp63^flox/flox^* recombination. These *de facto* LumI ΔΝp63**^+/-^** would still be able to contribute to clonal expansions since p63 heterozygous epithelia usually has no overt phenotype (Botchkarev and Flores, 2014). Additional clonal regulators that are independent of p63 might also play a role in LumI clonal growth.

**Figure 4.**
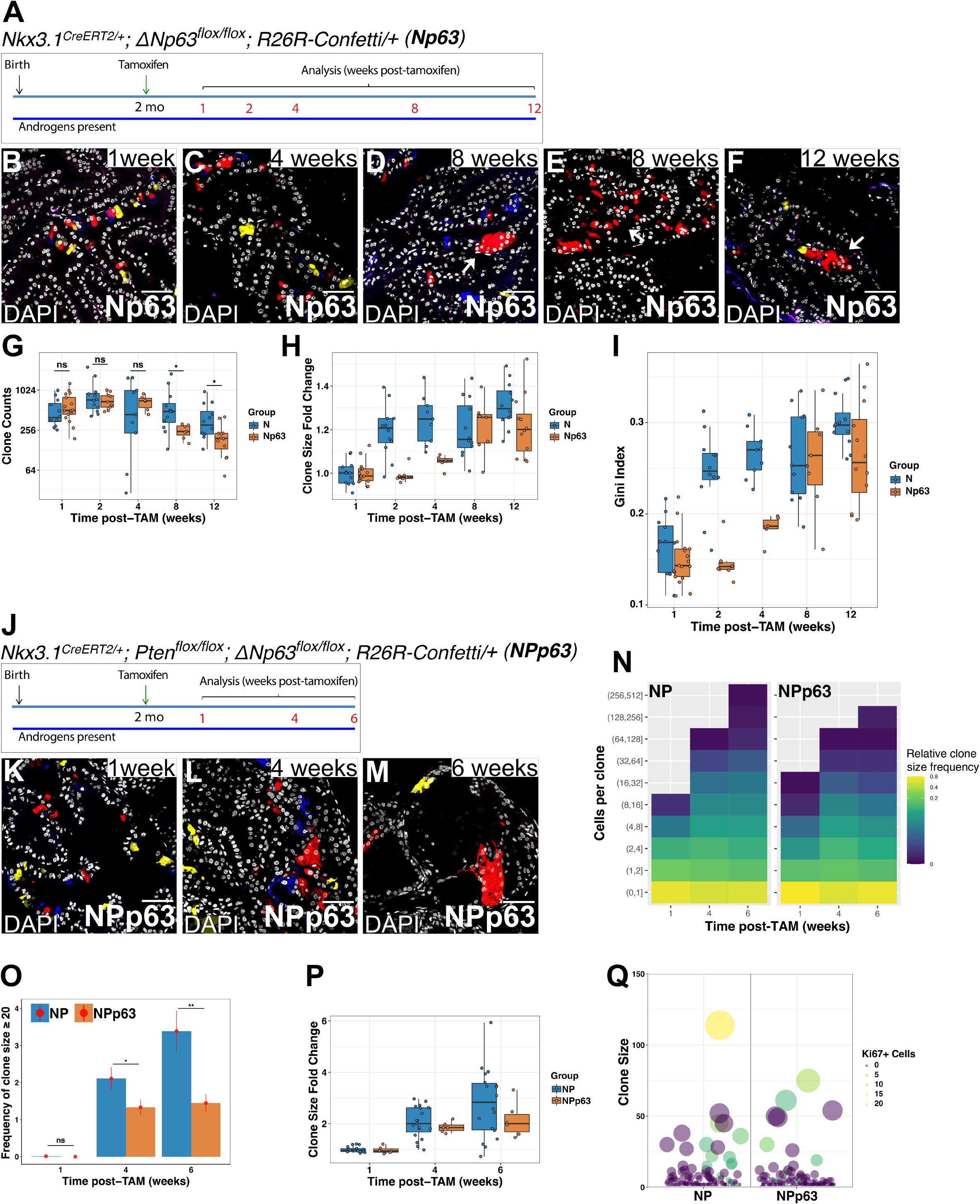
Genetic ablation of p63 in the luminal layer impairs clonal growth in normal prostate and PCa. **(A)** Time course of lineage-tracing experiments with the Confetti reporter in *Nkx3.1^CreERT2/+^; ΔNp63^flox/flox^* mice (*Np63*). (**B-F**) Representative images of clonal distribution in the Np63 anterior prostate at 1, 4, 8 and 12 weeks post-TAM, 40x images are shown. White arrows point to clones with heterogenous sizes in late timepoints. Scale bars represent 50 μm. (**G**) Clonal quantification of tile scan images of entire prostate sections showing a reduction in the total clones/section at 8 and 12 weeks in Np63 mice comparing to the N mice. (**H**) Clonal size fold change at each time point showing an impaired clonal expansion rate in Np63 mice at earlier timepoints. (**I**) Gini index of clonal sizes of *N-Confetti* mice and *Np63* mice showing reduced Gini index in the *Np63* mice at early timepoints indicating that inequality due to larger clones appears at a slower pace in *Np63* prostates. (**J**) Time course of lineage-tracing experiments with the Confetti reporter with *Nkx3.1^CreERT2/+^; Pten ^flox/flox^; ΔNp63^flox/flox^*mice (*NPp63*). (**K-M**) Representative images of clonal distribution in the *NPp63-Confetti* anterior prostate at 1, 4, 6 weeks post-TAM, 40x images are shown. White arrows point to clones with heterogenous sizes in late timepoints. Scale bars represent 50 μm. (**N**) Heatmap representation of clonal size distribution comparing densities of clones of various sizes at the indicated time points post-TAM in *NP* and *NPp63* tumors. While *NP* tumors maintain similar clones of medium size and a progressive increase in larger sized clones, the *NPp63* landscape accumulates fewer larger clones by 6 weeks. For *NP*: 4 mice per time point, 5-7 tile scans of entire sections/mouse, 37454 clonal events across all time points were included. For *NPp63*: 2 mice/per time point, 3 tile scans of entire sections/mouse, 7646 clonal events across all time points were included. Frequency is computed as the proportion of clones in a given size interval among all clones collected for a certain time point. A clone was defined as a contiguous fluorescent patch, including clones of 1 cell. (**O**) Frequency of larger clones (with more than 20 cells) in *NP* and *NPp63* tumors. (**P**) Confetti clonal size fold change in *NP* and *NPp63* tumors. On average, the clones grow slower in the *NPp63* tumors. (**Q**) Bubble plot of clone size vs their Ki67 cell status in *NP* and *NPp63* tumors showing fewer actively proliferating cells per clone upon p63 deletion in luminal cells. 5-6 regions of interest from *NP* and *NPp63* tumors were assessed. 430 clones from *NP* and 221 clones from *NPp63* were analyzed for their clone size and number of Ki67 positive cells in clones.

In sum, our *in vivo* lineage tracing of ***Np63*** mice validates ΔΝp63 as an important factor in maintenance of luminal clonal dynamics and demonstrates that factors affecting the LumI state can also alter the homeostatic clonal landscape.

To modulate the LumI in *cancer* cells, we generated ***NPp63*** mice which simultaneously delete *Pten* and *ΔΝp63* in Nkx3.1-Confetti cells and performed similar lineage tracing and clonal data acquisition (**Fig. 4J**). The ***NPp63*** tumors had similar histology to the *NP* tumors (**Suppl. Fig. 4E**). At 1 and 4 weeks of tracing, ***NPp63*** exhibited the same multiclonality as ***NP*** tumors (**Fig. 4K, 4L**). However, by 6 weeks, the ***NPp63*** clones appeared sparser with only few clones continuing to expand (**Fig. 4M**). To see if the remaining clones are completely p63 null, we performed p63 immunofluorescence analysis of the ***NPp63*** clones at 6 weeks post-tamoxifen. Most clones were p63-negative (**Suppl. Fig. 4G-J**), while few clones still contained p63-positive cells (**Suppl. Fig. 4K-N**), likely indicating heterozygous recombination of the p63^flox/flox^ alleles. Of note, most ***NP*** clones contained p63**^+^** cells (**Fig. 3O, Suppl. Fig. 4O**). Systematic quantification of ***NPp63*** clones further demonstrated the impaired clonal growth resulting from p63 genetic deletion in luminal cells. At 1 week post-induction, we observed a similar representation of clonal size distribution (**Fig. 4N)**. However, clonal growth patterns began to differ at 4 and 6 weeks. While ***NP*** tumors maintain similar clones of medium size and a progressive increase in larger sized clones, the ***NPp63*** landscape accumulates fewer larger clones by 6 weeks (**Fig. 4N**). In accord, larger clones (≥ 20 cells) had reduced frequency at 4 and 6 weeks in the ***NPp63*** tumors (**Fig. 4O**). Fold changes in the average clonal size in ***NPp63*** tumors were also reduced compared to ***NP*** tumors (**Fig. 4P**), indicating a general reduction of clonal expansion activities in the ***NPp63*** tumors. Staining for Ki67 paralleled the overall clonal growth patterns, with fewer actively proliferating cells per clone upon p63 deletion in luminal cells **(Fig. 4Q)**.

In sum, the tumor clones in ***NPp63*** are still viable but the overall expanding rate has been reduced. The few large clones maintained in the ***NPp63*** tumors are likely derived from cells with heterozygous recombination or unsuccessful recombination of the floxed *ΔΝp63* alleles. In addition, the other expanding clusters (e.g., Krt4**^+^**) might also be able to generate larger clones in accord with previous studies (Guo et al., 2020).

### The LumI features are enriched in mouse and human aging prostate and human prostate cancer

To understand the implications of our mouse findings for human prostate, we assessed the presence of “intermediate” cells by CK5/CK8 staining in adult/middle age and aging human normal prostate as well as in prostate cancer samples with various Gleason scores. Adult glands showed a normal distribution of basal (CK5**^+^**) and luminal (CK8**^+^**) layers (**Fig. 5A**). Even when the luminal layer is more abundant, the luminal positioned cells maintain a luminal phenotype (**Fig. 5A,** asterisk). However, the intermediate cells significantly increase in aging samples (**Fig. 5B, 5E**) and take over the glands in human prostate cancer samples (**Fig. 5C-E**), especially in those with higher Gleason score (**Fig. 5D)**.

**Figure 5.**
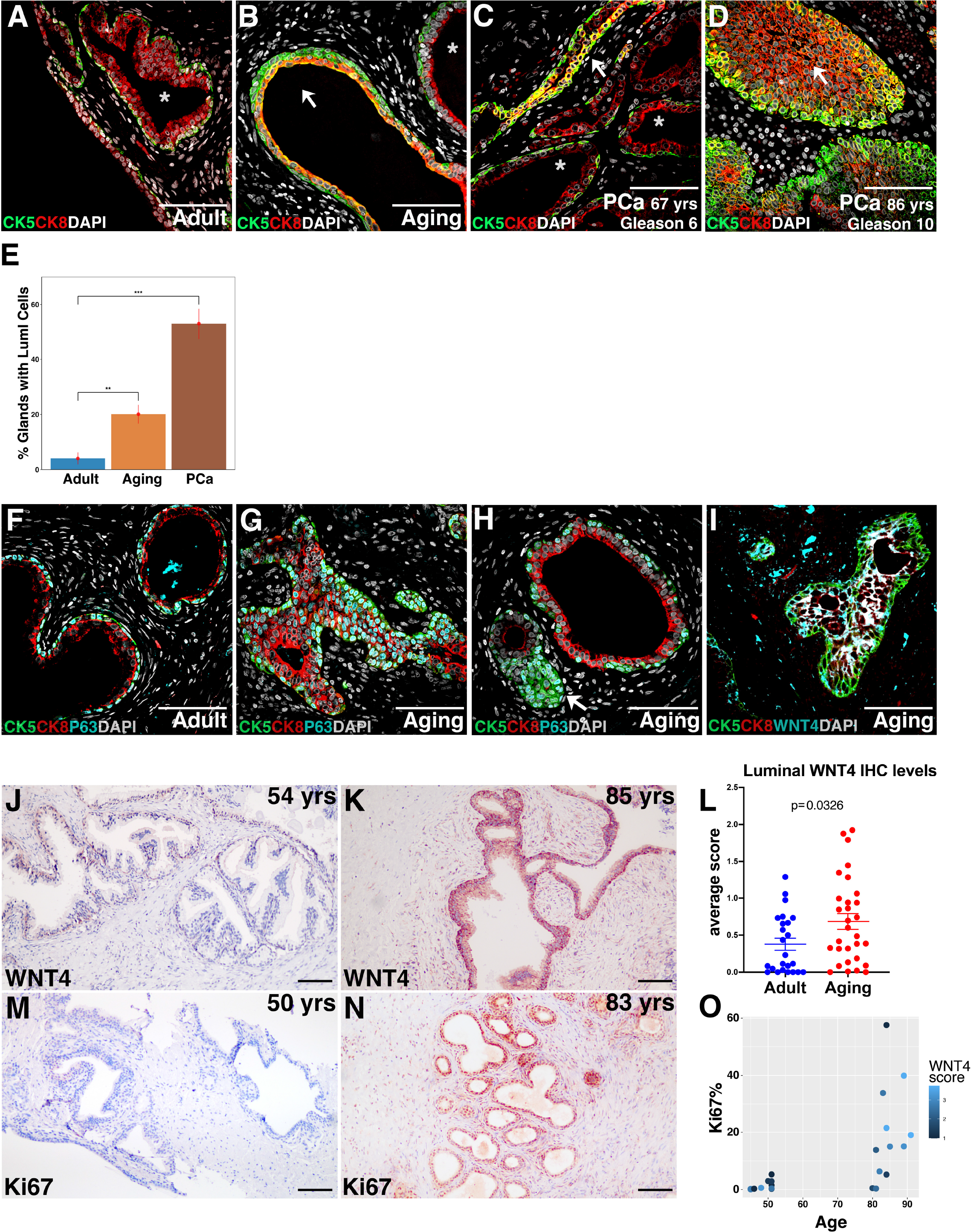
Intermediate cells are increasingly present with aging and cancer in human prostate. **(A-D)** “Intermediate” cells in human normal prostate samples (adult and aging) and PCa with various Gleason scores as shown by CK5/CK8 colocalization. (**E**) Distribution of glands with positive “Intermediate” cells (co-localization of CK5/CK8 in the luminal layer) in adult, aging and PCa samples. (**F**) Adult normal glands express p63 in all basal cells. (**G, H**) Aging normal glands exhibit p63 in luminal cells (**G**) along with glands that show increased basal cells (**H**). (**I**) Aged glands also express WNT4 in “luminal intermediate” cells. (**J-K**) Representative WNT4 immunohistochemistry staining in human adult (**J**) and aging (**K**) normal prostate gland. WNT4 is predominantly basal in adult normal prostate glands and increasingly luminal with aging. (**L**) An increase proportion of aging glands expresses WNT4 in both basal and luminal layer. A WNT4 score was given based on % positive cells and intensity of luminal Wnt4 staining. 24 Adult/middle age (49+/-3) and 30 Aging (85+/-3) human normal prostate samples were analyzed. 10-15 sections/sample were analyzed. (**M-N**) Representative Ki67 immunohistochemistry staining in human adult (M) and aging (N) normal prostate gland. Ki67**^+^** cells were rare in the adult group, while numerous glands in aging tissues express Ki67 in both basal and luminal layer. (**O**) The proportion of Ki67**^+^** cells increases with aging and correlates with the WNT4 score. Ki67 staining of 10 sections from the Adult/ middle age (49 +/- 3 years of age) group and 12 sections from the Aging (85 +/- 3 years of age) human normal prostate samples analyzed. Nuclei with positive Ki67 staining were quantified with ImageJ and normalized by total nuclei in each section.

In accord with our mouse data, the intermediate cell state is also associated with increased WNT4 and p63 levels (**Fig. 5F-I**). Specifically, most adult normal glands express p63 in all basal cells but not in luminal cells (**Fig. 5F**) while aging normal glands exhibit p63 in luminal cells (**Fig. 5G**) distinct from the basal proliferative glands that express p63 solely in the basal cells (**Fig. 5H**). Aging glands also express WNT4 in “luminal intermediate” cells (**Fig. 5I**). These analyses suggest that “luminal intermediate” glands co-exist with hyperproliferative basal glands in human aging tissue. Moreover, the “luminal intermediate” glands exist in human prostate cancer and aging prostate and might contribute to a hyperproliferative program. To quantify and further understand the correlation between WNT4 expression in aging prostate and cell proliferation we evaluated by IHC an additional 24 Adult/middle age (49 +/- 3 years of age) and 30 Aging (85 +/- 3 years of age) human *normal* prostate samples for WNT4 and Ki67. We found WNT4 to be predominantly basal in adult normal prostate glands (**Fig. 5J**). As expected from the IF analysis (**Fig. 5I**), an increase porportion of aging glands express WNT4 in both basal and luminal layer (**Fig. 5K**). For a quantitative assesment, we gave each sample a WNT4 score based on percentage (%) WNT4 positive luminal cells and the intensity of luminal WNT4 staining. Our analysis showed that the WNT4 luminal expression is significantly increased in the aging cohort (**Fig. 5L**, p=0.0326). Ki67+ cells were rare in the adult group (**Fig. 5M**); while numerous glands in aging tissues expressed Ki67 in both basal and luminal layer (**Fig. 5N**). By matching the Ki67 status with the WNT4 score for each sample, we found that the proportion of Ki67+ cells increases with aging and correlates with the WNT4 score (**Fig. 5O**). Taken together, our data suggests that the LumI intermediate cell state marked by unique Wnt4 and p63 expression might play an important role in aging prostate and prostate cancer.

## Discussion

We described here heterogeneity in clonal formation and clonal activities of prostate luminal cells, in which a majority of mature luminal cells remain quiescent while a small subset of cells proliferates and grows into clones to maintain the homeostasis. We uncovered a “luminal intermediate” transcriptional state (**LumI**) marked by unique Wnt/p63 signaling. The LumI state expands during long-term lineage tracing of the homeostatic prostate and constitutes a great portion of the cancer cell populations expanding in early stages of tumorigenesis. Moreover, the **LumI** cell state dramatically increases in human aging prostate samples and human prostate cancer samples. We also investigated the functional role of LumI cells in prostate homeostasis and prostate cancer growth *in vivo* by specific deletion of *ΔNp63*, a master regulator of this cells state, in luminal cells. ΔNp63 deficiency in LumI cells slowed clonal formation in homeostasis and reduced cancer clone size in prostate cancer models. Together, these results suggest that a luminal intermediate transcriptional state is functional in maintaining the tissue structure and homeostasis of androgen-intact prostate. Deregulations of luminal cells found in this intermediate transcriptional state may lead to prostate hyperproliferative diseases in aging including prostate cancer. Targeting this cell state, such as through loss of ΔNp63, may reduce the clonal formation activities in aging and cancer.

By combining scRNAseq analysis and lineage tracing of Nkx3.1**^+^**luminal cells, our analysis provides a *dynamic* perspective of the luminal heterogeneity in prostate homeostasis and prostate cancer. Recent studies of prostate luminal cells have also proposed a luminal progenitor cell type that is frequently double positive for CK5 and CK8 similar to LumI cells, but also expresses high PSCA and CK19 (Tang, 2021). Based on these markers, this cell type likely corresponds to our Krt4**^+^** cluster (PSCA high) defined in this work and previous studies (Guo et al., 2020). Indeed, in the integrated scRNAseq analysis with early prostate tumors, we also observed an increase in the Krt4**^+^** cell population, suggesting that the Krt4**^+^**cell type contributes to clonal growth in PCa. We also identified a progenitor-like cell cluster expressing marker genes including Tcastd2 and Ly6d, which mainly expands in the prostate cancer models and appears to be connected with the Krt4**^+^** cells. Our findings suggest that the LumI cell state we report here and the Krt4**^+^** cell types are two distinct cell states in the normal prostate luminal cell compartment. Recently, a study of a Pb-Cre driven Pten-deficient mouse model revealed an expansion of luminal intermediate cells occurring only upon malignant transformation (Germanos et al., 2022). Furthermore, underscoring the importance of these cells, the luminal intermediate phenotype expands post-castration in this model. Given that Pb-Cre promoter is expressed broadly in the epithelial cells of the prostate, the authors suggest that these cells might be attributed to multiple sources of origin: basal cells, luminal progenitors and luminal differentiated cells (Germanos et al., 2022). The Cre driver of our mouse models is restricted to the luminal compartment, thus, our findings encompass a *luminal-derived* luminal intermediate cell state.

We also demonstrated that loss of luminal ΔNp63 reduced clonal formation *in vivo*. In accord, a recent study using a lung adenocarcinoma (LUAD) mouse models demonstrated that *ΔNp63* promotes tumor initiation and progression: comparing *Kras^LSLG12D/+^* mice which developed lung adenomas and LUAD by 20 weeks, *ΔNp63^flox/flox^*; *Kras^LSLG12D/+^* (***Kp63***) mice showed no scorable adenomas or LUAD formation (Napoli et al., 2022). Moreover, using ChIP-Seq, the authors identified a ΔNp63 target gene, Bcl9l (member of Wnt family), as a key mediator of the oncogenic activities of ΔNp63 (Napoli et al., 2022). Of note, Wnt4 has been shown to represent a downstream target of p63 as well (Osada et al., 2006).

With potential translational implications, our integrated normal prostate – PCa scRNAseq analyses revealed the potential contribution of the LumI state to epithelial to mesenchymal transition (EMT). Within the EMT cell cluster 40% of the cells are CK5/CK8 double positive, suggesting that the intermediate cell state is also contributing to the EMT regulated cellular plasticity. The EMT cell cluster also shares the activated Wnt signaling profile and P63 regulon with the intermediate cell clusters. This is in line with previous research that has also demonstrated that activation of Wnt/β-catenin signaling correlates with the EMT phenotype of PCa cell lines (Jiang et al., 2007). Downregulating the Wnt/ β-catenin signaling pathway with specific inhibitors suppresses the EMT cell behavior in prostate cancer cells (Lee et al., 2019; Li et al., 2018; Sun et al., 2018). ΔNp63 has been recently shown to coordinate a transcriptional program that generates cancer stem cells residing in an EMT-like state in a breast cancer model (Lambert et al., 2022). This ΔNp63-dependent transcriptional program has been found to be distinct from the one that acts in basal mammary stem cells (Lambert et al., 2022). Our data also suggest that the ΔNp63- dependent LumI program is distinct from the role of ΔNp63 in basal prostate cells.

Altogether, we provide here a comprehensive view of the clonal dynamics of prostate luminal cells. A small subset of the prostate luminal cells is able to form larger clones in prostate homeostasis. Single cell RNA-seq revealed an intermediate cell state that is expanding in the course of homeostatic lineage tracing and also contributes to prostate cancer formation. Understanding how this intermediate cell state is regulated in homeostasis and deregulated in hyperproliferative contexts is crucial for the development of prognostic markers and new therapeutic targets of prostate diseases.

## Methods

### Mouse strains and genotyping

All experiments involving animals were performed according to protocols approved by the Institutional Animal Care and Use Committee at Stony Brook University. The *Nkx3.1^CreERT2/+^* (Wang et al., 2009), *Pten^flox/flox^* (Groszer et al., 2001), *K-Ras^LSLG12D/+^* (Jackson et al., 2001), *ΔNp63^flox/flox^* (Chakravarti et al., 2014) (kind gift from Elsa R. Flores, Moffitt Cancer Center), and *R26R-Confetti* (Snippert et al., 2010) targeted alleles have been described previously. The compound mice of the desired phenotype, *Nkx3.1^CreERT2^; R26r-Confetti (**N**)* and *Nkx3.1^CreERT2/+;^ Pten^flox/flox^; R26r-Confetti (**NP**), Nkx3.1^CreERT2/+^; Pten^flox/flox^; K-Ras^LSL-G12D/+^; R26r-Confetti (**NPK**); Nkx3.1^CreERT2/+;^* ΔNp63*^flox/flox^; R26r-Confetti (**Np63**)* and *Nkx3.1^CreERT2/+;^ Pten^flox/flox^*; ΔNp63*^flox/flox^; R26r-Confetti (**NPp63**)* were obtained by crossing mice carrying these alleles. All mice were maintained in C57BL/6N background. All studies were done using littermates that were genotyped prior to enrollment; mice were randomly enrolled to time groups and only male mice were used because of the focus on prostate.

Genotyping was performed by PCR using tail genomic DNA, with the following primers (full sequences are listed in **Suppl. Table 5** - reagents) for the following alleles: *Nkx3.1* wild-type, *CreERT2* and its alternative CreTK, *Pten^flox^*, *R26R–Confetti*, *K-Ras^LSL-G12D^*, and ΔNp63*^flox^*.

### Mouse procedures

For lineage-marking and simultaneous deletion of Pten and/or activation of Kras*^G12D^* mutation in Nkx3.1 expressing prostate cells, *N, NP* and *NPK* mice were administered tamoxifen (Sigma; 10.8 mg/40 g body weight in corn oil) by oral gavage once daily for 1 day at the age of two months, followed by various chase periods of 1, 2, 4, 6, 8 and 12 weeks. For lineage-marking and simultaneous deletion of ΔNp63 and Pten, *Np63* and *NPp63* mice were administered tamoxifen in the same way. Middle aged *N* mice were administered one dose of tamoxifen at the age of 12-14 months. For collecting the lineage traced cells for scRNAseq, the *N, NP*, and *NPK* mice were administered tamoxifen (Sigma; 10.8 mg/40 g body weight in corn oil) by oral gavage once daily for 4 days at the age of two months.

### Tissue collection

For histological, confocal imaging and immunofluorescence analysis, individual prostate lobes were dissected and fixed in 4% paraformaldehyde for 1-2 hours for subsequent cryopreservation in OCT compound (Sakura). Prostate lobes were also fixed in 10% formalin followed by paraffin embedding and sectioning for H&E and IHC staining.

### Confocal imaging and clonal quantification

Prostate tissues (AP lobes) were systematically sectioned through the entire volume as 10 and 20 μm sections. 3 slides with 6 sections of 20 μm were followed by one slide with 6 sections at 10 μm, then 6 sections of 20 μm were discarded and this sectioning plan was repeated throughout the whole embedded tissue. From each of the slide with 20 μm sections, one section was used for direct visualization of the Confetti colors by confocal imaging or tile scanning. Slides were washed with phosphate buffered saline (PBS) followed by staining of nuclei with Hoechst 33342 (Invitrogen) and mounting with SlowFade Gold Antifade reagent. For a precise quantification of clusters/prostate tumor, we acquired images of entire prostate (AP lobes) sections by automated tile scanning with a Leica SP8X confocal and merging of the acquired “tiles” using the LAS-X software. The slides with 10 μm sections were used for immunofluorescence staining.

To quantify the Confetti clones present in the tile images of prostate sections we developed an automated image processing script based on previous methods for quantification of immunofluorescence images (Lee et al., 2014) and the Image Processing Toolbox from Matlab (MathWorks, Inc.). The program reads the raw confocal file, performs median filtering to remove the noise, subtracts background followed by thresholding and binarization, hole filling, removing of small objects, detecting individual regions of fluorescence in each channel, and counting contiguous fluorescent cluster, and counting Hoechst**^+^** nuclei. The program extracts the total number of color clusters/channel, the total number of clusters/image and the total cell number/each cluster.

### Histology and immunostaining

Hematoxylin–eosin (H&E) staining was performed using standard protocols on 3 μm paraffin sections. Histological assessments were performed using a published classification of mouse PIN lesions (Park et al., 2002).

For immunofluorescence, 10 μm cryosections were subjected to permeabilization with 0.5% Triton/PBS (nuclear antigens) or 0.05% Tween-20/PBS (cytoplasmic and membranous antigens) for 30 minutes, blocking in 5% goat serum/PBS for 1 hour followed by incubation with primary antibodies (see antibody list in **Suppl. Table 5** - reagents) at 4°C overnight in humidified chambers. Alexa Fluor 633 or 647 (Life Technologies) were used for secondary antibodies to distinguish the immunofluorescence signal from the endogenous fluorescence of the Confetti fluorochromes. Harsh retrieval steps were omitted to avoid destruction of the Confetti expression. Fluorescence images were acquired using a Leica TCS SP8X spectral confocal microscope (Stony Brook University) with Leica LAS-X software.

For immunohistochemistry staining, 3μm paraffin sections were. We used primary antibodies (see **Suppl. Table 1**). The primary antibodies were recognized by SignalStain Boost IHC Detection Reagents (HRP, secondary antibodies mouse and rabbit, Cell Signaling) and visualized by VECTOR NovaRED Peroxidase Substrate Kit (Vector Laboratories). The positive staining signals of WNT4 in luminal cells of human samples were assessed by two individuals independently. Ki67**^+^** cells were quantified using ImageJ software.

### Confocal imaging and Ki67 quantification in clones

To quantify proliferating populations within clones, 10 μm cryosections were immunostained with Ki67 primary antibody and Far Red secondary antibody as described above. Sections were imaged on the Leica TCS SP8X by identifying regions of interest (ROIs) containing multiple and diverse clones. Using LASX software, multiple 4x4 40x tile scans (total 16 tiles) were acquired of various ROIs throughout the section and merged to create representative images of each ROI. Images were processed and quantified via Matlab as described above. Each clone detected by Matlab was assigned a number for identification purposes. Next, Ki67^+^ cells were hand counted for each clone in each ROI. Using the clonal identification numbers, Ki67 counts were matched to each clone in the image.

### Tissue dissociation and flowcytometry

Tissue dissociation and isolation were performed as previously described (Chua et al., 2014). Briefly, mouse anterior (AP) prostate lobes were dissected in cold phosphate buffered saline (PBS) and minced with scissors followed by incubation in DMEM/F12 (Gibco) supplemented with 5% FBS and 1:10 dilution of collagenase/hyaluronidase (STEMCELL Technologies) at 37°C for 3 hr. Dissociated tissues were spun at 350 g for 5 min, and resuspended in TrypLE (Gibco), followed by incubation at 37°C for 20 minutes. After centrifugation at 350 g, pelleted cells were resuspended with pre-warmed 5 mg/ml dispase (STEMCELL Technologies) supplemented with 1:10 dilution of 1 mg/ml DNase I (STEMCELL Technologies), triturated vigorously for 2 min, and diluted by addition of HBSS/2% FBS. Finally, the cell suspension was passed through a 40 μm cell strainer (Falcon). Filtered single cells were resuspended in Opti-MEM (Gibco) and flow sorted based on RFP, CFP or YFP expression on a BD FACS Aria II instrument. GFP was underrepresented as previously reported (Snippert et al., 2010).

### Single-cell RNA sequencing

For cells used for Single-cell RNA sequencing (scRNAseq), cells from prostate AP lobes were FACS sorted as described. For each time point and each scRNAseq sample, cells from AP lobes from 3-4 mice/genotype were pooled prior to FACS sorting. Sorted cells were washed and resuspended at a concentration of 800 cells/μl. Single-cell libraries were constructed with 10x Genomics Chromium Single Cell 3′ Library & Gel Bead Kit v3. The Confetti**^+^** cells were processed for loading on a Chromium Next GEM Chip for 10X Chromium Controller processing. In total, 20 μl of cell suspension was loaded per sample aiming for a cell recovery of 10,000 cells. Single-cell capture, reverse transcription, cDNA amplification and 3’ gene expression library construction were performed according to Chromium Single Cell 3**ʹ** Reagent Kits v3 user guide. Sequencing was performed with an Illumina Novaseq 4000 instrument (Novogene).

### Computational analysis Processing of scRNAseq data

The Cell Ranger Single-cell analysis pipelines (*cellranger 3.0.2*) were used to process the FASTQ files generated from sequencing. First, we used the cellranger count pipeline to perform alignment with the 10x Genomics pre-built mouse genome for mm10-3.0.0 followed by filtering, barcode counting, and UMI counting in default parameter setting. From this, we obtained reads of 7533 cells from the *N* sample at 1 week of tracing and 5512 cells from the *N* sample at 8 weeks of tracing; 7048 cells from the *NP* mice and 15923 cells from the *NPK* mice. The gene expression matrix for each library from Cell Ranger output was then loaded to RStudio for subsequent pre- processing. We used R package *Seurat* (version 3.0) for quality control filtering. First, the expression matrix was trimmed for each gene that was expressed in at least three cells and each cell has at least 200 genes. Then, only cells with a number of unique features between 500 to 4100 were kept to avoid low-quality cells, empty droplets or cell doublets. In addition, the percentage of reads that map to the mitochondrial genome was capped at 15% to exclude low-quality/dying cells which often exhibit extensive mitochondrial contamination.

### Sample integration and cluster identification using *Seurat*

For sample integration and cluster identification, we continued to utilize the *Seura*t pipeline. First, the preprocessed Seurat objects representing the samples of 1 week and 8 weeks cells were normalized with default setting. Next, we used the function *FindVariableFeatures* to calculate a subset of features that exhibits high cell-to-cell variation in the dataset. Then we used the *FindIntegrationAnchors* function to identify anchors for integrating the two datasets from the two timepoints samples. This step was followed by the standard *Seurat* workflow for visualization on the integrated object with default settings. Specifically, we used the ‘*Scaledata’* function to scale and center features in the dataset and the ‘*RunPCA*’ function to perform principal component analysis (PCA) to reduce dimensionality of the dataset to the top 30 PCs. followed by Uniform Manifold Approximation and Projection (UMAP) dimensional reduction on the first 12 PCs. The Seurat function *FindNeighbors* was used to construct a Shared Nearest Neighbor (SNN) graph based on k-nearest neighbors of each cell. Then clusters of cells were identified with the ‘*FindClusters*’ function with a resolution of 0.5.

Using Seurat functions such as *FindAllMarkers*, *VlnPlot* and *DimPlot*, we identified and visualized marker genes for each respective cluster. Dying cells with high mitochondrial gene expression and other non-epithelial cells were eliminated from downstream analyses. After characterization of all cell clusters, some clusters that appear very similar based on marker gene expression were manually merged. Some major clusters were subjected to sub-clustering analysis where sub-clusters were extracted from the original Seurat object with the ‘*subset*’ function.

We performed a similar *Seurat*-based integration analysis of cells from *N*, *NP* and *NPK* mice to draw comparisons between normal and malignant epithelial cells.

### Single cell trajectory analysis with *Monocle*

The N/NP/NPK integrated scRNAseq data from Seurat was also used for trajectory analysis with *Monocle*. First, the *DDRTree* dimensionality reduction was applied with default parameters. Next, we used the *orderCells* function to learn the trajectory from the reduced space embeddings. To order the cells within the trajectory, we selected ordering the genes with the unsupervised procedure called “*dpFeature*” which extracted genes that differ between clusters.

### Gene regulatory network analysis using *SCENIC*

Cluster specific gene regulatory networks were identified with *SCENIC* R package and *SCENIC* pipeline as previously described (Aibar et al., 2017). The *SCENIC* algorithm makes use of the *AUCell* algorithm to identify and score the activity of regulons in each cluster by analyzing gene expression of known transcription factors and their targets. The results were plotted using *ComplexHeatmap* (Gu et al., 2016).

### Enrichment analysis with Gene Set Variation Analysis (GSVA) and *clusterProfiler*

Gene signatures from individual clusters of the N/NP/NPK integrated object were enriched using GSVA function (Hänzelmann et al., 2013). Pathways to be examined were collected from the Molecular Signatures Database (MSigDB) collections, specifically the Hallmark gene sets and the curated Kyoto Encyclopedia of Genes and Genomes (KEGG) gene sets. To compare GO enrichment changes between the 1 week sample and the 8 weeks sample, we used the *enrichGO* function of *clusterProfiler* (Wu et al., 2021; Yu et al., 2012) on a set of upregulated genes from the Intermediate1/2 clusters of the 8 weeks sample and visualized the enriched GO categories with the built-in *dotplot* function.

### Statistical analysis

R language and GraphPad Prism (GraphPad Software, Inc) were used as main tools for statistical analysis. A p-value <0.05 was considered to be statistically significant. Statistical analyses were performed using a two-sample *t*-test, or Wilcoxon Rank Sum test as appropriate. Details of each experiment were explained in figure legends. Data in results are reported as mean *±* SEM unless otherwise stated. The index of dispersion (σ^2^ /μ) was used to access the diversity of clone sizes.

### Data availability

The scRNAseq data of 1 week and 8 weeks post-tamoxifen N prostate cells are deposited in the Gene Expression Omnibus (GEO) database under accession number GSE216764. The NPK scRNAseq generated for this study are deposited under accession number GSE196371.

### Code availability

All codes used in this manuscript are publicly available. A list of the R packages used is provided in **Suppl. Table 6**.

## Acknowledgements

We would like to acknowledge the technical support provided by the Research Histology Core Laboratory - Department of Pathology - Stony Brook Medicine and Stony Brook Cancer Center. The authors wish to acknowledge the Stony Brook Medicine Biobank/Stony Brook Cancer Center Tissue Analytics Shared Resource for identifying clinical cases and procuring the FFPE blocks from the Pathology clinical archive collection. We are indebted to Dr. Michael Shen for the Nkx3.1^CreERT2^ mouse models and to Dr. Elsa Flores for the ΔΝp63^flox/flox^ mice. We thank Todd Rueb for technical support in flow cytometry. We thank Dr. Scott Powers for access to the 10x Genomics Chromium Controller and Dr. Manisha Rao for assistance with the 10x Genomics protocol. This work was supported by 1 F99 DK131481-01 to F.L., NIGMS 1R35GM122561-01, Stony Brook Cancer Center Pilot Funds and the Laufer Center for Physical and Quantitative Biology to G.B., 5K22CA188169 and 5R21CA270610 to F.T., Elsa Pardee Foundation Award to F.T., lab start-up funds provided by the Department of Urology and Stony Brook Cancer Center to F.T.

## Author contributions

F.T and F.L. designed the study, F.L., L.F.T., S.R., D.R., K.T., J.G.R., A.N. and F.T. performed the experiments, F.L. and F.T. performed the computational analyses, M.S. and G.B. developed the automated image quantification, J.L. performed the histopathology assessment, G.B. and A.N. provided review and editing of the manuscript, F.T. and F.L. interpreted the data and wrote the manuscript.

## Competing interests

The authors declare no competing interests.

## Supplemental Figure Legends

**Supplemental Figure 1.**
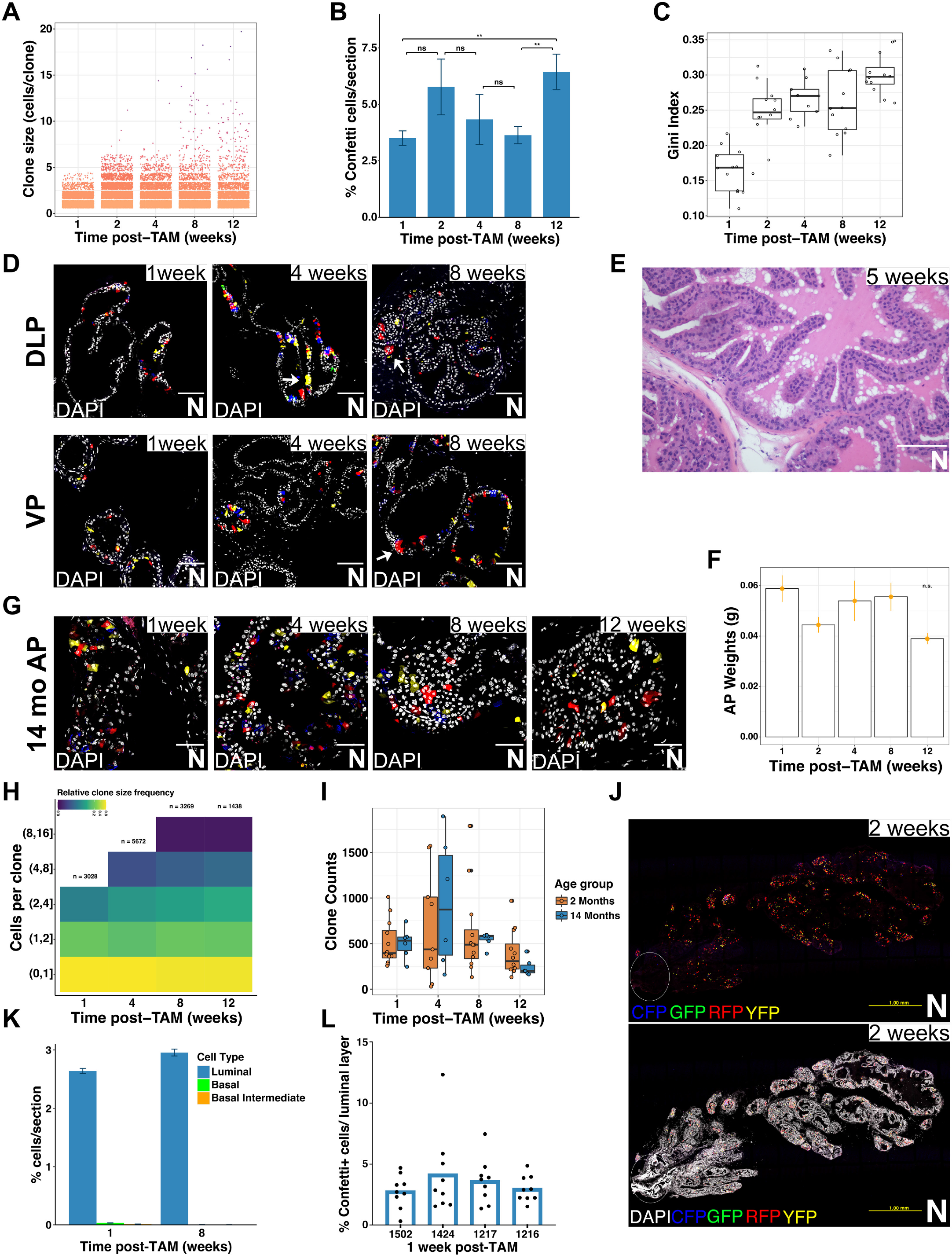
Analysis of clonal dynamics in *N* mice. (**A**) Clonal size distribution for each time point in the AP lobes. Each dot represents a clone. 3 sections of each AP lobe per mouse are included. 4-5 mice per time point. (**B**) Quantification of Confetti**^+^** cells for each time point in *N* mice anterior prostate (AP) lobes. The numbers were extracted from the MatLab image quantification script as described in the methods section. Mean and SEM of total Confetti**^+^** cells/total number of nuclei/section are shown. (**C**) The Gini index (Gini coefficient) of clone sizes calculated using the R package *ineq*. The Gini coefficient measures the inequality of the clonal size distribution. The higher Gini index at later timepoints suggests increased heterogeneity of clonal sizes. (**D**) Representative images of clonal distribution in the *N- Confetti* dorsolateral (DLP) and ventral (VP) prostate lobes at 1, 4 and 8 weeks post-TAM. The same increase in the number of larger clones with the time of tracing as in the AP lobes is observed. 20x images are shown, scale bars represent 100 μm. (**E**) H&E staining confirmed normal structure in *N* prostate AP lobes by 5 weeks post-induction. Scale bar represents 100 μm. (**F**) Weights of AP lobes collected at each time point. Constant weights support the fact that no aberrant growth occurred during the lineage tracing. (**G**) Representative images of clonal distribution in anterior prostates of middle-aged *N-Confetti* mice at 1, 4, 8 and 12 weeks post-TAM, 40x images are shown, scale bars represent 50 μm. (**H**) Heatmap representation of clonal size distribution comparing densities of clones of various sizes at indicated time points post-TAM in middle-aged *N-Confetti* mice. 3 mice per time point, 3 tile scans of entire sections/mouse, 24360 clonal events across all time points were included. A clone was defined as a contiguous fluorescent patch, including clones of 1 cell. (**I**) Comparison of clone counts in adult (2 Months) and middle-aged (14 Months) *N-Confetti* mice. The clone counts at each time point remains stable in the middle-aged group and larger clones become more abundant at later time points similar to adult homeostasis dynamics. (**J**) Representative scan of Confetti clonal distribution in *N-Confetti* AP lobes at 2 weeks post-TAM. An entire section of an AP lobe is shown as a 20x scan. Confetti signals are shown on the top panel and Confetti signals overlaying with DAPI signals are shown on the bottom panel. The periurethral/proximal prostate region of the AP is on the left side of the scan (circle). Note the absence of Confetti clones in this region. (**K**) Positional and histologic analysis of the proportion of Confetti- labeled cells among the main prostate cell types in 1 week and 8 weeks *N* samples. 9 20x images from 3 sections of each mouse were examined for labeled cell types, 4 mice of each time point were used. (**L**) Quantification of Confetti**^+^** cells in the luminal layer at 1 week post-TAM in *N* mice AP lobes. The luminal cells were manually counted. Data points are sampled from 9 20x images per mouse, 4 mice were included.

**Supplemental Figure 2.**
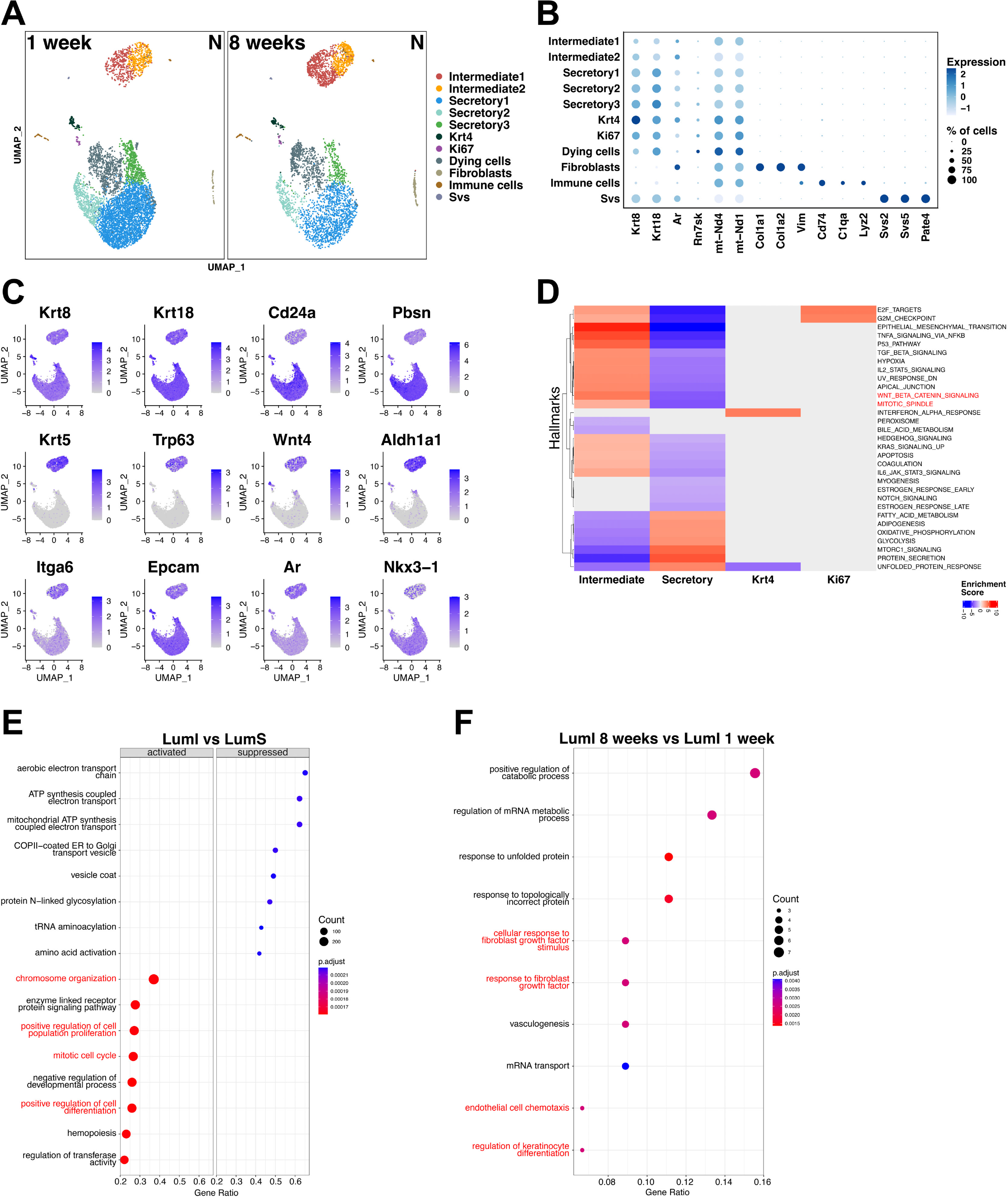
scRNAseq of Nkx3.1-lineage traced cells reveals cellular heterogeneity within the luminal layer. **(A)** Raw cell type cluster distribution in 1 week and 8 weeks post-TAM *N* samples by UMAP (Seurat) identifying a total of 11 cell clusters in both samples. The small non-epithelial, non-prostatic or dying cells clusters were discarded in subsequent analyses. (**B**) Dot plot of selected marker genes in all the original clusters. (**C**) Feature plot of selected marker genes expressed in LumI, LumS or both. (**D**) Heatmap of Hallmark geneset expression enrichment in LumI, LumS, Krt4**^+^** and Ki67**^+^**clusters. For simplicity, the Intermediate1 and Intermediate2 cells were integrated in one Intermediate cluster, while the Secretory1, 2, and 3 were integrated in one Secretory cluster. (**E**) Pathway enrichment analysis with *clusterProfiler* in LumI vs LumS clusters. (**F**) Pathway enrichment analysis with *clusterProfiler* in LumI clusters from 1 week vs LumI clusters from 8 weeks.

**Supplemental Figure 3.**
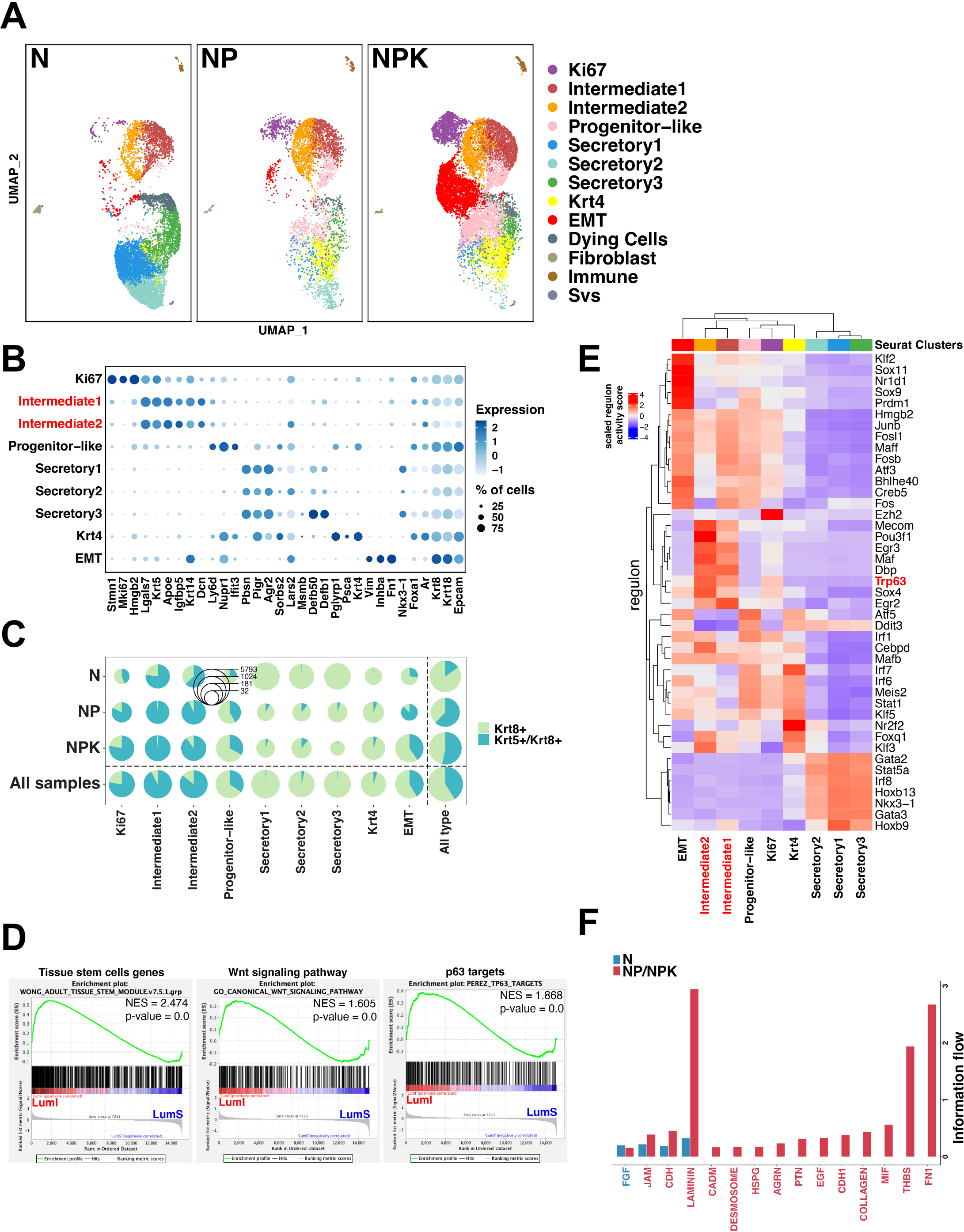
Comparative analysis of scRNAseq of Nkx3.1-lineage traced cells in *N*, *NP* and *NPK* mouse models. **(A**) Raw cell type cluster distribution of integrated *N*, *NP* and *NPK* samples by UMAP (Seurat) identifying a total of 13 cell clusters in all samples. The small non-epithelial clusters contaminating the FACS sorting were discarded in subsequent *Seurat* analyses. (**B**) Dot plot of selected marker genes in epithelial clusters. Note that transcriptomic features of the *N* Intermediate cell clusters are maintained in the cancer samples. (**C**) The proportion of Ck5+/Ck8+ cells increases in cancer. (**D**) GSEA of adult tissue stem cell markers (left), canonical Wnt signaling pathway (middle) and p63 targets (right) indicate enrichment of these three gene sets in the malignant LumI cell state. (**E**) Regulon analysis by *SCENIC* of the N/NP/NPK combined *Seurat* object indicates that p63 is maintained as a master regulator of the intermediate clusters in all samples and likely plays an important coordinating role in the expansion of cancer clones. (**F**) Signaling pathway networks that are strongly active in LumI cells of the *N* samples or LumI cells of the *NP/NPK* samples based on the differences of overall information flow as predicted by *CellChat*. The overall information flow of a signaling network is calculated by summarizing all the communication probabilities in that network.

**Supplemental Figure 4.**
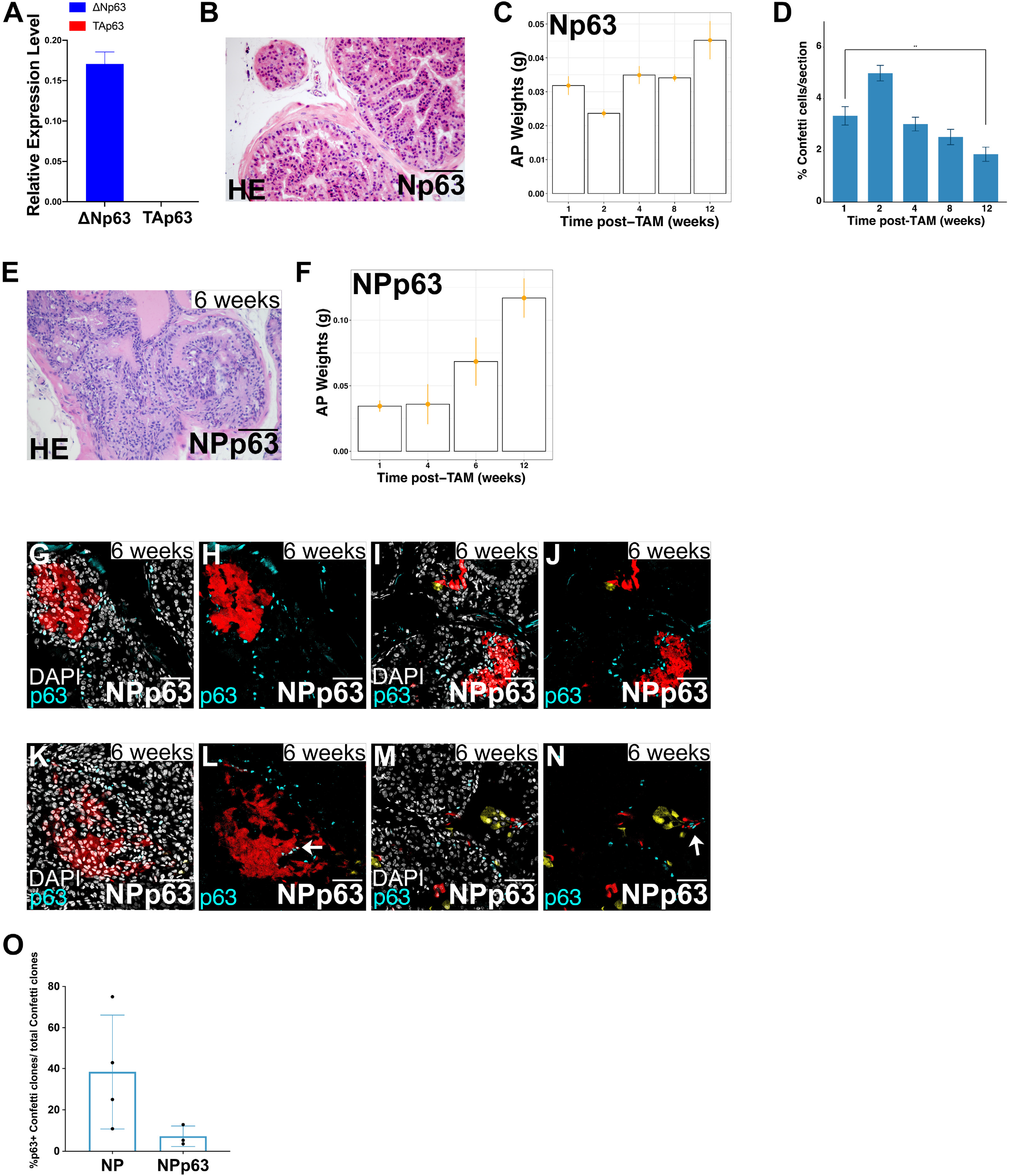
Characterization of the *Np63* and *NPp63* mouse models. **(A)** Relative expression levels of the main p63 isoforms in Confetti^+^ cells of N prostate. Note that the ΔΝp63 isoform is expressed at much higher levels than TAp63 in prostate luminal cells. **(B)** H&E staining of an *Np63* anterior prostate at 1 week post-TAM showing normal tissue structure. Scale bar represents 100 μm. (**C**) Weights of Np63 AP lobes collected at each time point. Constant weight indicates that no aberrant growth occurred during lineage tracing. (**D**) Quantification of Confetti**^+^** cells for each time point in Np63 mice AP lobes. Mean and SEM of total Confetti**^+^**cells/total number of nuclei/section shown. (**E**) Representative H&E staining of *NPp63* tumors at 6 weeks post-TAM showing features of low-grade PIN similar to *NP* mice. (**F**) Weights of *NPp63* AP lobes collected at each time point. Increased weight reflects the aberrant growth occurring during *NPp63* transformation upon tamoxifen administration. (**G-J**) Most of the *NPp63* Confetti clones had no detectable p63 expression. (**K-N**) Rare Confetti clones with p63 expression in *NPp63* mice likely indicating clones with heterozygous recombination of the inducible *p63^flox/flox^* alleles. (**O**) Quantification of p63**^+^** Confetti clones in *NP* and *NPp63* AP lobes. Clones with p63**^+^** cells are lost in the NPp63 lobes.

## Notes

### Competing Interest Statement

The authors have declared no competing interest.

